# Connectomic consistency: a systematic stability analysis of structural and functional connectivity

**DOI:** 10.1101/862730

**Authors:** Yusuf Osmanlıoğlu, Jacob A. Alappatt, Drew Parker, Ragini Verma

## Abstract

Connectomics, the study of brain connectivity, has become an indispensable tool in neuroscientific research as it provides insights into brain organization. Connectomes are generated for different modalities such as using diffusion MRI to capture structural organization of the brain or using functional MRI to elaborate brain’s functional organization. Understanding links between structural and functional organizations is crucial in explaining how observed behavior emerges from the underlying neurobiological mechanisms. Many studies have investigated how these two organizations relate to each other; however, we still lack a proper understanding on how much variation should be expected on the two modalities, both between people cross-sectionally and within a single person longitudinally. Notably, connectomes of both modalities were shown to have significant differences depending on how they are generated. In this study, for both modalities, we systematically analyzed consistency of connectomes, that is the similarity between connectomes in terms of individual connections between brain regions or in terms of overall network topology. We present a comprehensive study of consistency in structural and resting-state functional connectomes both for a single subject examined longitudinally and across a large cohort of subjects cross-sectionally. We compared connectomes generated by different tracking algorithms, parcellations, edge weighting schemes, and edge pruning techniques. We evaluated consistency both at the levels of individual edges using correlation and at the level of network topology via graph matching accuracy. We also examined consistency of connectomes that are generated using most commonly applied communication schemes. Our results demonstrate varying degrees of consistency for the two modalities, with structural connectomes showing higher consistency than functional connectomes. Moreover, we observed a wide variation in consistency depending on how connectomes are generated. Our study sets a reference point for consistency of connectome types, which is especially important for structure-function coupling studies in evaluating mismatches between modalities.

## 1. Introduction

Within the last two decades, connectomics, the study of brain connectivity, has emerged as an effective way of analyzing structural and functional connectivity of the brain where the brain network is modeled as a graph with nodes representing the brain regions and weighted edges representing the strength of connectivity between region pairs [Sporns et al., 2005]. Structural connectivity is obtained using fiber tracking of diffusion MRI (dMRI) to estimate the macroscopic axon fiber bundles that connect pairs of brain regions [Hagmann et al. 2008]. Functional connectivity of the brain, on the other hand, is generally calculated as the temporal correlation between the activation of brain regions, which is commonly obtained using resting state functional MRI (rs-fMRI) [Fox et al., 2007]. Understanding the relationships between structural and functional connectivity of the brain is essential to explain how human behavior emerges from the neurobiological organism [Menon et al., 2011].

Analysis of connectomes have brought to light several significant neuroscientific findings regarding the network organisation of the structural [van den Heuvel et al., 2013] and functional [Greicius et al., 2003] connectivity of the brain, as well as how the two organisation types relate to each other [Osmanlıoğlu et al., 2019]. Compounded with the fact that both dMRI and resting state-fMRI (rs-fMRI) have their own inherent limitations [Buckner et al., 2013, Schilling et al., 2018], such as noise and the indirect relation between the measurements and the ensuing model [Assaf et al., 2019], connectomes of both modalities were shown to have significant differences depending on how they are generated [Kelly et al., 2012, Sotiropoulos and Zalesky, 2019]. In generation of structural connectomes from dMRI data, for example, the two commonly used tracking approaches (deterministic or probabilistic tracking) are known to result in connectomes of varying densities and organizations [Bonilha et al., 2015]. Since such differences might potentially be a source of conflicting results in connectomics studies, a proper understanding of how much variation should be expected on both structural and functional connectivities is necessary. Such an analysis is especially important for structure-function coupling analysis, which assumes that functional activations emerge from the structural pathways connecting brain regions [Honey et al., 2009], since discrepancies observed between the two modalities might be emerging from the inherent variations of the connectivity types.

Recent years have seen several studies evaluating the consistency of structural [Owen et al., 2013] and functional [Guo et al., 2012, Horien et al., 2019] connectomes that are generated using differing schemes. Consistency, in this context, is defined as the variation of the connectivity measures across connectomes [Moussa et al., 2012]. Although consistency is evaluated previously for structure and function of the brain separately, comparison of consistency between the two modalities on the same dataset is still lacking. In order to alleviate this need for a comprehensive assessment of variation in connectomes as a result of various factors, we present an extensive consistency analysis of structural and resting state functional connectomes that are derived from dMRI and rs-fMRI data, respectively.

In our analysis, we evaluated consistency of connectomes both for a single subject examined longitudinally and across a large cohort of subjects cross-sectionally. We assessed consistency in both structural and functional connectomes across three parcellation schemes and various thresholds for pruning weak edges. In further analysis of structural consistency, we compared structural connectomes generated by probabilistic and deterministic tracking algorithms through various edge weighting schemes. In investigating consistency for functional connectomes further, we evaluated positive, negative, and whole functional connectivities separately. We evaluated consistency both at the levels of individual edges using correlation and at the level of network topology via graph matching accuracy. We also examined consistency of connectomes that are generated using shortest path and weighted communicability, two commonly applied communication schemes in the study of brain network organisation.

## 2. Data and Methods

### 2.1. Datasets

In this study, the publicly available MyConnectome and Philadelphia Neurodevelopmental Cohort (PNC) datasets were used for evaluating the consistency of structural and resting state functional brain connectivity data, where the first dataset facilitated investigating within-subject consistency across time while the second dataset provided insights on the between-subject consistency cross-sectionally.

MyConnectome data contains 106 sessions of diffusion weighted MRI (dMRI), resting-state functional MRI (rs-fMRI), and task fMRI scans of a 45 years old healthy male recorded within an 18 month period [Poldrack et al., 2015]. In this study, data from 16 sessions acquired within a 5 month period with both dMRI and rs-fMRI data acquired on the same day, were used. All of the structural and functional imaging data used in this study was acquired on the same Siemens 3T Skyra scanner using a 32-channel head coil. A single anatomical T1-weighted image was acquired using an MP-RAGE sequence with TR/TE = 2400/2.14 ms at an isotropic 0.8 mm resolution. The rsfMRI data were collected using a multi-band Stejskal–Tanner EPI sequence (TR/TE = 1160/30 ms, flip angle = 63 degrees) with a voxel size of 2.4⨉2.4⨉2.0 mm across 512 volumes obtained in a 10 minute scan. The dMRI data were acquired using a multi-band Stejskal–Tanner EPI sequence, with opposite gradient readout options (L>R, and L<R) in order to enable distortion correction. The data were acquired at high resolution of 1.7 mm isotropic voxel size with a TR/TE = 5000/108 ms, in 60 directions with two shells (b=1,000 s/mm^2^ and 2,000 s/mm^2^) of 30 directions each and four b=0 volumes.

The PNC dataset consists of dMRI and rs-fMRI scans of 1445 unique subjects aged 8-22 years [Satterthwaite et al., 2014]. For our analyses, participants with poor structural and functional imaging data quality or a history that suggested potential abnormalities of brain development were excluded. In this study, the rs-fMRI and dMRI images of 694 healthy subjects aged 8-22 years (mean=15.71, SD=±3.28 years), including 280 males and 414 females, were considered. All data were acquired on the same Siemens 3T Verio scanner, using a 32-channel head coil. For each subject, a T1-weighted anatomical scan was acquired, along with the rs-fMRI and dMRI scans. The T1-weighted MP-RAGE sequence was acquired at a resolution of 1 mm isotropic voxel size with a TR/TE = 1810/3.5 ms. The rs-fMRI data were acquired with 3mm isotropic voxels at a TR/TE = 3000/32 ms, for 124 volumes where the scan time was 6 min 18 sec. The dMRI data were acquired with a 2mm isotropic voxel size, at a TR/TE = 8100/82 ms in 64 diffusion directions with a b = 1000 s/mm^2^ and seven b = 0 volumes.

### 2.2. Preprocessing

#### T1 preprocessing

Every T1 image in the PNC dataset and the single anatomical T1 image from the MyConnectome Project underwent bias correction using the N4BiasCorrection tool from ANTs [Tustison et al., 2010] followed by skull stripping using Muse [Doshi et al., 2016].

#### Resting state - functional MRI preprocessing

In order to calculate functional connectomes, both the PNC and MyConnectome rs-fMRI data were first preprocessed for motion correction [Woolrich et al., 2001]. The first 6 volumes of the motion-corrected data were subsequently removed, and the timeseries were slice-time corrected and co-registered to their corresponding processed anatomical MP-RAGE image. Spikes in intensity across volumes were then estimated, volumes with excessive spiking were excluded from further analysis, and signal obtained from voxels containing non-gray matter tissue were regressed out.

#### Diffusion MRI preprocessing

To calculate the structural connectomes, the dMRI data from both datasets were denoised [Manjón et al., 2013], corrected for motion and eddy current distortion [Andersson et al., 2016], bias-corrected [Tustison et al., 2010], and finally skull stripped. MyConnectome data were further corrected for EPI distortion as the data were acquired with opposite gradient readout options [Smith et al., 2004]. A standard tensor model was then fit to the single-shell PNC data, and the b=1000 shell of the MyConnectome data using DIPY to calculate fractional anisotropy (FA) maps, which were used for registrations [Garyfallidis et al., 2014].

### 2.3. Atlases and Registrations

In order to evaluate the effect of parcellation on connectomic consistency, structural and functional connectomes were calculated at three parcellation scales of the Schaefer atlas [Schaefer et al., 2018]. The three parcellations, hereafter referred to as Schaefer 100, Schaefer 200, and Schaefer 300 based on the number of regions, were defined in MNI space and had to be registered to each subject.

For each dataset, the T1 image was used as the target of registration for both the labels defined in MNI space, and the data defined in dMRI space. Using a deformable registration from ANTs [Avants et al., 2007], we computed warps from template space (MNI) to a given subjects’ T1 image. Additionally, registrations were computed between a subject’s T1 image and the processed FA map, using two pairs of fixed and moving images (T1-FA, and T1-B0). It is important to note that in the MyConnectome dataset, having only a single anatomical T1 image, all 16 dMRI sessions were registered to this T1, which was also the target of registration from MNI space. Using this registration pipeline, labels defined in MNI space were brought to diffusion and anatomical T1 spaces.

### 2.4. Connectomes

#### Resting-state Functional Connectomes

After rigid registration of the preprocessed BOLD data to the T1 [Jenkinson et al., 2002], timeseries were extracted using the atlases defined in the T1 space. Pearson’s correlation was then calculated between timeseries of each pair of ROIs and Fisher’s z-transform was applied to the resulting matrices to obtain functional connectomes.

#### Structural Connectomes

In order to evaluate the impact of different tracking algorithms on connectomic consistency, deterministic (SD_STREAM) and probabilistic (IFOD-2) tractography were separately used to calculate connectomes [Tournier et al., 2012, Tournier et al., 2010]. Both methods were run using Anatomically Constrained Tractography (ACT)[Smith et al., 2012] in mrtrix3. The tractography options (min/max tract lengths = 5/400 mm, step size = 1, angle = 60°, and streamline count = 500K) were identical for both tracking methods. Seeding for tractography across both algorithms was performed at the gray matter-white matter interface. Base structural connectomes were generated by designating the connectivity between pairs of ROIs as the total number of streamlines between them. To evaluate the effect of scaling on consistency, two further connectomes were derived from the base connectome by i) log scaling the streamline counts and ii) scaling each streamline count by the combined volume of its ROI pair.

### 2.5. Consistency Measures

Connectomic consistency is defined as the variation of the connectivity measures across connectomes. Given a measure that quantifies the similarity between two connectomes, we calculated consistency over a dataset as follows. We first calculated the similarity between a subject and the rest of the dataset, yielding a similarity score for each. We then rendered the average of these values as the similarity score of the query sample relative to the rest of the dataset. We finally considered the mean of these similarity scores as a measure of connectomic consistency, where higher values indicate that the generated connectomes are consistent across the dataset.

Since connectomic consistency is inherently contingent upon the choice of similarity measure, we evaluated similarity using two measures that highlight different aspects of connectivity in a brain network. We first considered *Pearson’s correlation* (denoted *r* in the text), which is commonly used to assess similarity between connectomes [Hagmann et al. 2008, Amico and Goni, 2018]. Representing values in the upper triangle of connectomes as a vector, we regarded Pearson’s correlation between the vector representation of two connectomes as their similarity, which is a value in [−1,1] with larger values indicating higher similarity.

As an alternative to correlation, we considered *matching accuracy* (denoted MA in the text) which we recently proposed as a similarity measure for connectomes [Osmanlıoğlu et al., 2019]. Matching accuracy is a measure driven from the solution to the graph matching problem, where the goal of matching is to find a mapping between the nodes of two given graphs by coupling nodes which resemble one another, while considering the overall connectivity structure of the network [Osmanlıoğlu and Shokoufandeh, 2015]. In order to achieve this goal, we formulated graph matching as a combinatorial optimization problem as follows: given two sets of nodes A and B, and a cost function *c*: A ⨉ B → ℝ determining the cost of assigning each node in A to a corresponding node in B, the aim is to calculate a one-to-one mapping *f*: A → B between the nodes of the two sets by minimizing 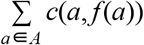. In the context of connectomics, we regarded the assignment cost between nodes as the Euclidean distance between the *k*-dimensional feature vectors of nodes which encode the connectivity of a node relative to the rest of the nodes in a parcellation with *k* ROIs. We solved the optimization problem by using the Hungarian algorithm [Kuhn, 1955]. In the resulting mapping, we regarded the percentage of nodes that were correctly mapped to their counterparts as the matching accuracy between the two connectomes, which is a value in [0,100] with larger values indicating higher similarity.

Although the two measures quantify the consistency of the networks encoded across connectomes, the two measures differ in their utilization of the network structure. Pearson’s correlation can be considered to be a measure that is sensitive to local similarity, as it compares corresponding edges between two connectomes while ignoring the topology of the network. On the other hand, graph matching accuracy quantifies the similarity of the network topology of two connectomes by considering the connectivity signatures of all regions while determining the optimal mapping between their nodes. Briefly, Pearson’s correlation quantifies the consistency of connections locally, whereas graph matching accuracy quantifies the consistency of the network topology.

### 2.6. Experimental Setup

In our analysis, we carried out a comprehensive set of experiments that can broadly be categorized into four groups.

1. We first compared the consistency of structural and functional connectomes. In our analysis of structural connectomes, we considered deterministic and probabilistic connectomes separately, while in functional connectomes, we evaluated the full functional connectome, as well as positive and negative functional connectivity by removing all negative and positive edges, respectively.
2. We then evaluated consistency of structural connectomes that are obtained through two common post-processing steps of i) scaling edges and ii) calculating “traffic patterns” [Goñi et al., 2014]. Among edge scaling schemes, we compared unscaled connectomes to those scaled by the log function and by node volume. As studies investigating structure-function coupling in the brain mainly rely on connectivity maps that are obtained by applying certain traffic schemes over the structural connectivity of brain regions, we compared the consistency of connectomes obtained through two prominent communication schemes to that of direct connectivity between regions. Being the most commonly applied traffic pattern in brain studies, the first communication scheme we evaluated was the weighted shortest path [Honey et al., 2009], which assumes that communication in the brain occurs through the shortest path between node pairs. To complement this scheme, we also considered weighted communicability [Estrada and Hatano, 2008] which assumes that communication occurs not only through the shortest path, but additionally through suboptimal routes.
3. We further evaluated the consistency of structural connectomes after removing the weakest edges to achieve various target network densities, a commonly applied post-processing method over structural connectomes to remove spurious fibers generated by tracking algorithms.
4. Finally, we evaluated the consistency of positive, negative, and full functional connectomes after removing edges lower than various thresholds, which is a common method to remove noise in functional connectivity maps.

Throughout our analyses, we investigated connectomic consistency at three parcellations of the Schaefer atlas (100, 200, and 300 ROIs), over both MyConnectome and PNC datasets, using both correlation and matching accuracy as the connectomic consistency measures.

## 3. Results

### 3.1. Comparison of consistency in structural and functional connectomes

In order to get an overall view of connectomic consistency, we first compared the consistency of structure and function on two datasets using the two similarity measures (Fig. 1). Our results highlighted a significantly higher consistency in structure relative to function across all testing conditions. Focusing on consistency across datasets, we observed that both structural and functional connectivity is more consistent in the longitudinal MyConnectome data relative to the cross-sectional PNC data. We further observed a reduced consistency both in structure and function as the parcellation gets finer across all test cases. Comparing tracking methods, we observed that probabilistic connectomes demonstrate higher consistency relative to deterministic connectomes. In comparing functional connectivity types, we observed that positive functional connectivity has higher consistency than full functional connectivity, where the difference is larger over PNC dataset relative to MyConnectome dataset. We also observed that, negative functional connectivity has the lowest consistency across all connectivity types with a large margin of difference. Finally, investigating the effect of similarity measures, we observed that both correlation and matching accuracy demonstrated the same consistency patterns in that, the ordering of consistency scores across experiments were identical among the measure types. Additionally, we noted a variation in the magnitude of consistency differences across modalities. We observed that matching accuracy demonstrates a smaller difference of consistency across structural connectivity types and positive and full connectivity, relative to correlation. In consistency of negative functional connectivity, however, matching accuracy demonstrated a larger consistency difference relative to other connectivity types, in contrast to correlation reporting a decent level of consistency.

**Figure 1.**
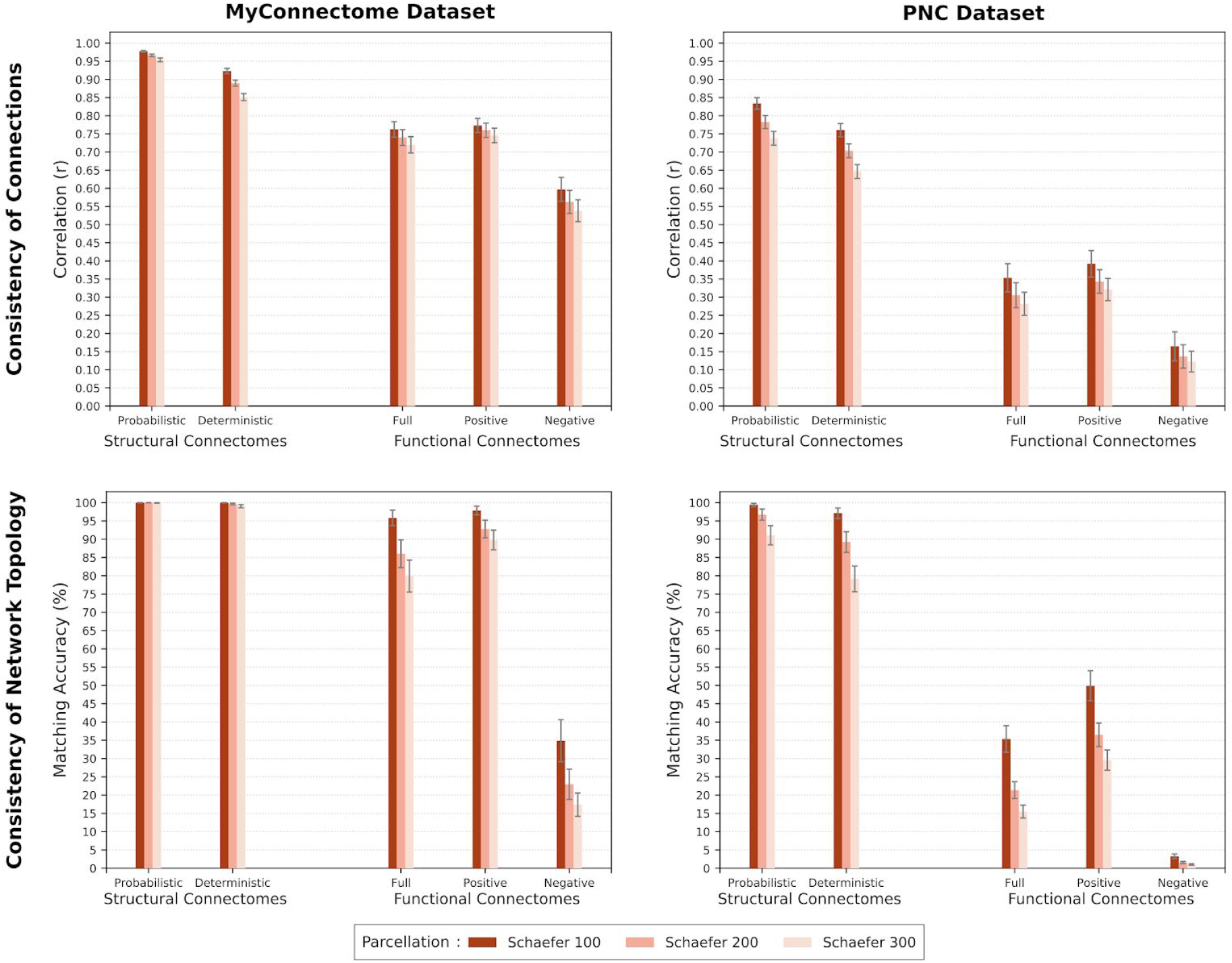
Comparison of structural and functional connectomic consistency. Top row shows correlation based consistency results which highlight stability at the level of individual connections, whereas bottom row shows matching accuracy based consistency results highlighting stability of the network topology. Left column demonstrates consistency of structure and function on the same subject over time while the right column demonstrate the consistency across a large healthy cohort. (Error bars indicate standard deviation in consistency scores.)

### 3.2. Consistency of structural connectomes

We investigated the consistency of structural connectivity in further detail (Fig. 2 and 3). We observed that connectomes generated through probabilistic tracking were more consistent than those generated through deterministic tracking across all testing conditions. In line with the results of the previous experiment (Section 3.1), we observed that consistency is lower at finer parcellations across both datasets, and higher across the MyConnectome dataset relative to the PNC dataset. Additionally, we noted that the standard deviation of consistency scores is larger in the PNC dataset relative to that of the MyConnectome dataset. Comparing consistency across communication patterns, we observed that connectomes obtained by applying the weighted communicability scheme over direct connectivity maps achieved highest consistency, followed by direct connections and finally the shortest path. However, we noted that the effect of communication patterns on consistency was larger in the PNC dataset compared to that of the MyConnectome dataset. When investigating the effect of scaling, we observed that log scaling of edges results in a higher consistency relative to no scaling in most of the cases, whereas the effect of scaling edges by node volumes on consistency is not significant in the majority of the test cases. We observed that, although matching accuracy based consistency was high and stable for the majority of the cases, shortest path based connectomes without log scaling demonstrated a relatively lower consistency, especially on the PNC dataset, for both of the tracking methods.

**Figure 2.**
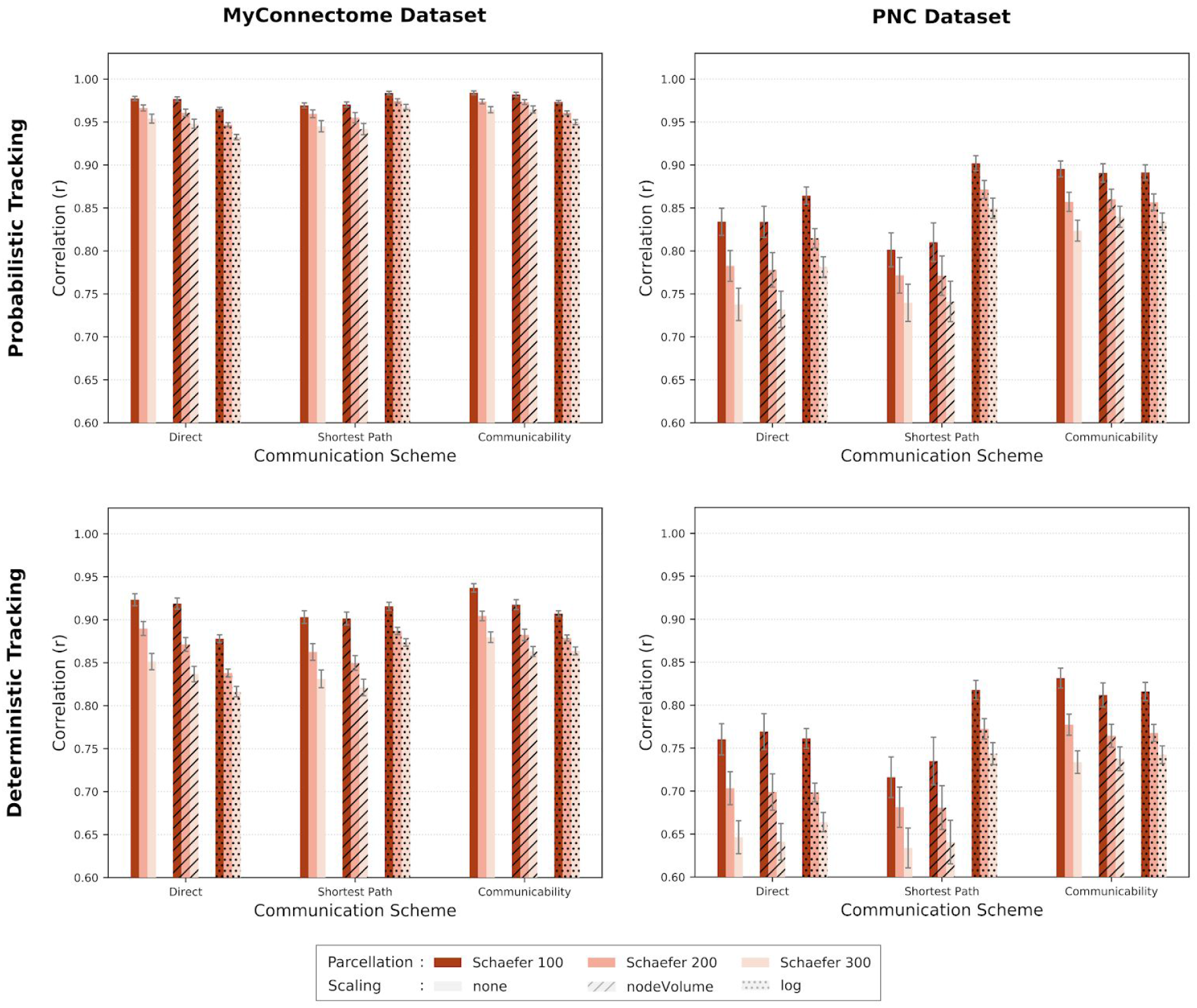
Consistency of connections in structural connectivity maps of deterministic and probabilistic tracking as quantified by Pearson’s correlation. Consistency is evaluated for two communication patterns in addition to direct connectivity, across two scaling schemes along with no scaling, over the two datasets. (Error bars indicate standard deviation in consistency scores. See Fig. 3 for consistency of network topology for the same testing conditions.)

**Figure 3.**
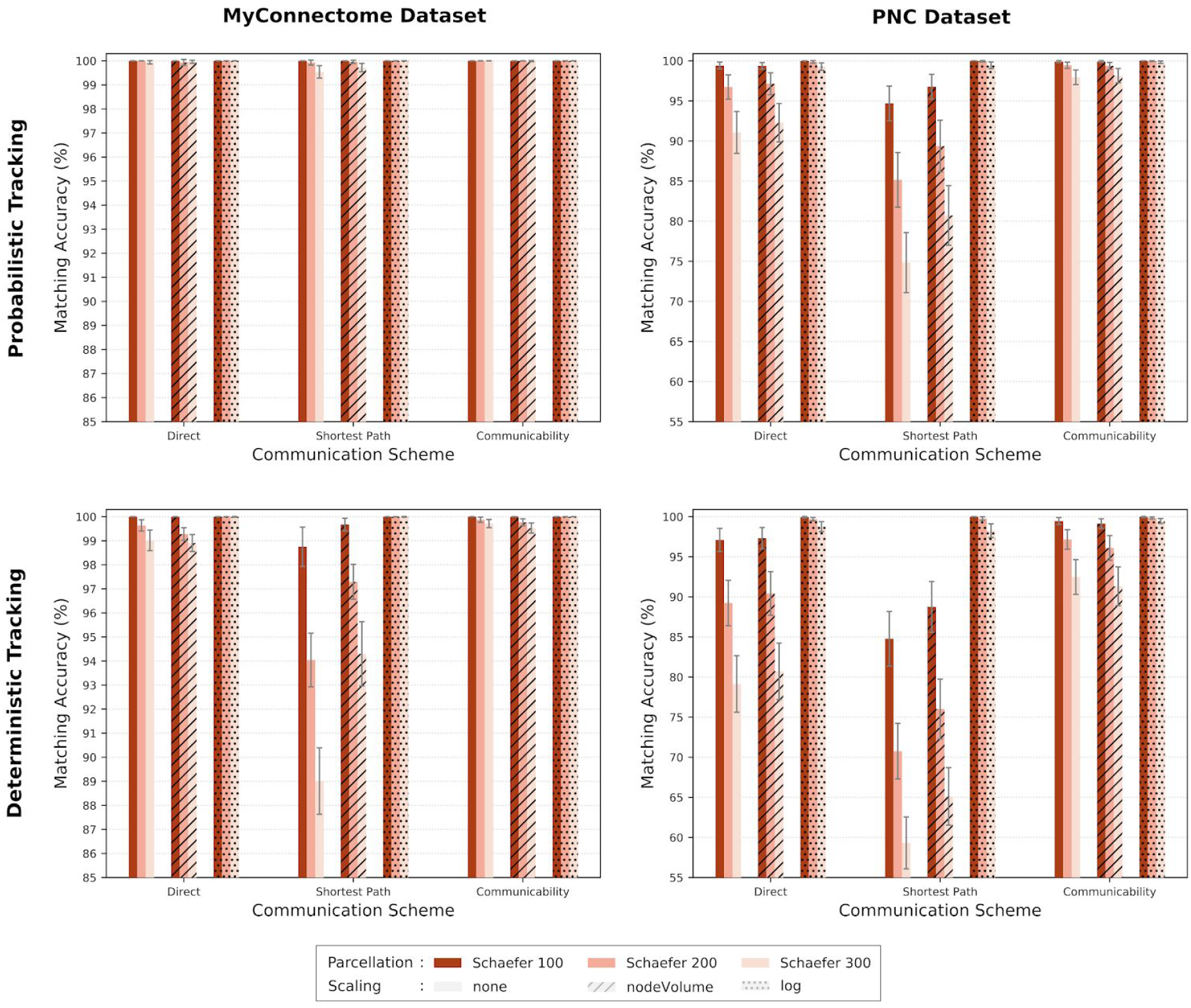
Consistency of network topology in structural connectivity maps of deterministic and probabilistic tracking as quantified by matching accuracy. Consistency is evaluated for two communication patterns in addition to direct connectivity, across two scaling schemes along with no scaling, over the two datasets. (Error bars indicate standard deviation in consistency scores. See Fig. 2 for consistency of connections for the same testing conditions.)

### 3.3. Consistency of deterministic and probabilistic structural connectomes with thresholding

Furthering our investigation on structural connectomes, we next evaluated the effect of thresholding on consistency across a range of densities, by removing lower weighted edges to set the nonzero edge density of the connectome at a certain level. In this experiment, we investigated direct connectivities (i.e., traffic patterns are not applied) without any scaling, over three parcellations, and across two datasets by using the two consistency measures. Noting that the mean density across connectomes were recorded as shown in Table 1 before any thresholding, we didn’t observe any significant change in consistency until the thresholding removed the majority of the edges which resulted in reduced consistency. To give a specific example, noting that densities of connectomes of MyConnectome and PNC datasets were (56.81%,52.94%) for probabilistic and (28.55%,29.32%) for deterministic tracking, we observed that consistency didn’t show any significant change over Schaefer 100 parcellation until the threshold of 10% density, while the consistency reduced rapidly with lowering density further. We also observed that this critical threshold value was lower over finer parcellations, which had a lower density before any thresholding, such as the Schaefer 300 parcellation where consistency didn’t change until the density is reduced below 5%. However, we noted that the structural consistency at the highly sparse density of 2% was still higher than the consistency of functional connectivity, such as the probabilistic structural consistency scores of (r=0.94, MA=83.5% on MyConnectome) and (r=0.67, MA=57% on PNC) in contrast to positive functional consistency scores of (r=0.77, MA=97% on MyConnectome) and (r=0.39, MA=50% on PNC) on Schaefer 100 parcellation. We further observed that, compared to correlation, matching accuracy based consistency was affected less by thresholding for higher densities while it demonstrates a steeper decay in consistency as the edges are removed further than the critical threshold. We also note that there were no significant differences between the consistency patterns of probabilistic and deterministic connectomes across thresholds.

**Table 1.** Ratio of non-zero edges to all possible number of edges in deterministic and probabilistic structural connectomes across the two datasets at three parcellations before thresholding. Densities are shown in percentages.

|  | MyConnectome |  | PNC |  |
| --- | --- | --- | --- | --- |
|  | Probabilistic | Deterministic | Probabilistic | Deterministic |
| **Schaefer 100** | 56.81 | 28.55 | 52.94 | 29.32 |
| **Schaefer 200** | 37.31 | 15.62 | 33.31 | 16.12 |
| **Schaefer 300** | 27.18 | 10.22 | 23.68 | 10.8 |

### 3.4. Consistency in functional connectomes

Finally, we detail our analysis on the consistency of functional connectomes (Fig. 5). In all test cases, we observe that full and positive connectomes achieve higher consistency relative to negative connectomes, with positive connectomes having a slightly higher accuracy than the full functional connectome. As in previous experiments, we observed that connectomes with coarser parcellations have higher consistency. We also observed that, removing edges with weights smaller than a magnitude of 0.2 did not have a significant effect on the consistency of full and positive connectomes while it reduced consistency of negative connectomes even for magnutes as small as 0.05. However, we noted a steady decrease in consistency over full and positive connectomes as well. Comparing the two datasets, we observed a lower consistency in all functional connectivity types over PNC. We also noted that when compared with correlation based consistency, matching accuracy based consistency reported a larger difference between the negative connectivity and the full and positive connectivities.

## 4. Discussion

### Structural connectivity is more consistent than functional connectivity

Throughout our analyses, the one common result that clearly stands out is the higher consistency of structural connectomes relative to functional connectomes (Fig. 1-5). Despite the fact that both dMRI and rs-fMRI modalities have their own inherent limitations [Buckner et al., 2013, Schilling et al., 2018], structural connectivity is expected to demonstrate higher consistency than functional connectivity due to the more dynamic nature of the latter [Chang and Glover, 2010]. Our comparative results indicate that the consistent structural network topology of the brain does not directly translate to a comparably consistent functional network structure.

Although consistency of both structural [Owen et al., 2013] and resting state functional [Guo et al., 2012, Horien et al., 2019] connectivities in the human brain were previously reported, to the best of our knowledge, ours is the first study that contrasts the consistency of dMRI and rs-fMRI over the same experimental setup. This result is important in providing a reference point for the connectomics studies especially that involve data in one of the modalities (for example, dMRI) and refer to the literature that report on the other modality (such as rs-fMRI) in order to interpret implications of their results. Thus, we recommend researchers to take the difference of consistency between the dMRI and rs-fMRI connectomes into account when making inferences across the modalities.

### Consistency differences have consequences in joint structure-function studies

The effect of the disparity in connectomic consistency across structural and resting state functional connectivity is especially acute in the structure-function coupling, where the goal is to find a mapping between the functional activations in brain and the underlying structural pathways connecting regions [Honey et al., 2009]. In network neuroscience, the brain is modelled as an information processing network [Bassett and Sporns, 2017] such that the structural pathways connecting brain regions act as conduits through which the information flows to generate functional interactions among brain regions. In such a setup, indirect functional interactions between regions are considered to happen by information exchange through intermediate regions, which is assumed to follow a certain communication scheme. Over the last decade, several communication schemes were suggested as the traffic pattern of the brain including shortest path [van den Heuvel et al., 2012], path transitivity, search information [Goñi et al., 2014], and communicability [Estrada and Hatano, 2008].

Despite several attempts at finding the communication scheme of the brain, structure-function coupling studies have reported decent correlations between the structural and functional connectomes (r≲0.5), indicating that the alignment between the structural connectivity and functional activations are far from perfect [Honey et al., 2009, Goñi et al., 2014, Osmanlıoğlu et al., 2019]. One common explanation for this is the inability of the devised communication schemes in capturing the complex functional interactions in brain. Results that we presented in this study might provide further insight into this problem. We showed that, although the structural connectomes have relatively small variation across the subjects, functional connectivities that are deemed to emerge from underlying structural pathways demonstrate a larger variety (Fig. 1). This might indicate that the functional data captured during the rs-fMRI scan is limited in describing the overall functional connectivity between brain regions.

We note that, the connectomes based on direct connections are generally sparse connected graphs, such as the ones we used in our experiments with 10% to 60% edge densities varying according to the tracking method and parcellation, whereas communicability and shortest path based connectomes are fully connected graphs since they represent strength of connectivity between any two node pairs which might be through direct or indirect connections. Our results in this study demonstrated that communicability based connectomes achieve highest consistency, which is followed by direct connection and shortest path based connectomes (Fig. 2-3). This result indicates that communicability based connectomes, which foresees communication occurring through multiple suboptimal paths in addition to the optimal shortest path, provides a connectivity map that has more commonalities across subjects, relative to that based on the shortest path, which restricts communication to occur exclusively through the optimal path. Although the shortest path has been the most commonly adopted scheme to model the communication in the brain in the connectomics literature, we recently showed that weighted communicability describes the functional connectivity of brain regions better than other known communication patterns. Our previous study also demonstrated that, direct connections are better than shortest path in explaining the functional connectivity of the brain [Osmanlıoğlu et al., 2019]. Thus, our results in this study demonstrating the same ordering among these three connectivity schemes add another dimension to the choice of communication pattern in structure-function coupling.

### Consistency within a person and between people are different

Connectomes have been shown to have aspects unique to the person leading to the emergence of the concept of connectome fingerprinting [Mars et al., 2018], which suggests the identifiability of a human by their brain connectivity pattern. This has been evaluated on structural [Osher et al., 2015] and functional [Passingham et al., 2002, Finn et al., 2015] connectomes, separately. These results have demonstrated that, two connectomes of a subject obtained from scans at different time points generally resemble each other more than connectomes of other subjects. Our results demonstrate a near perfect consistency for the structural connectivity and a very high consistency for the functional connectivity on a subset of the MyConnectome dataset, which consists of repeated scans of the same subject within a 5 months period. This highlights the identity preserving aspect of the connectomic fingerprinting for a single subject (Fig. 1). Comparing our results on the MyConnectome with the PNC, we observed that consistency across different subjects is lower both in structure and function. This indicates the presence of unique connectivity patterns of subjects which highlights the identity differentiation ability of connectomes, leading to a connectomic fingerprinting across subjects. However, this latter result also points to a common connectivity backbone across subjects in both structure and function, as we still observe decent consistency levels in both modalities.

We observed a larger difference on functional than structural connectomic consistency between MyConnectome relative to PNC, which might be due to a combination of reasons. Firstly, the longer rs-fMRI acquisition time of the MyConnectome data, which is 1.5 times longer than that of the PNC data, might have provided a more complete picture of the functional connectivity in a subject. Secondly, compared to the structure, functional connectivity might contain more aspects unique to the person that can be used as a connectivity fingerprint. Finally, lower consistency across subjects in functional connectivity might be due to the activation map being parcellated with a common atlas across all subjects when generating the connectomes, as it was recently stated that functional parcel boundaries reconfigure with cognitive state [Salehi et al., 2019].

### Processing of the data affects consistency

With the exponential increase in the number of scientific publications [Bornmann and Mutz, 2015], reproducibility of research findings has become a major problem [Ioannidis, 2005]. Neuroimaging research and connectomics, as rapidly growing fields of science along with the complicated methods that are used to process and evaluate data [Bandettini, 2019], faces unique reproducibility challenges [Picciotto, 2018, Zuo et al., 2019]. A major source of concern in connectomics in this regard stems from the complicated preprocessing pipelines applied over raw imaging data (such as motion correction, denoising, and registration), involved methods to process curated data (such as tractography on structure and independent component analysis on function), and additional post-processing steps employed before obtaining finalized connectomes (such as thresholding or scaling of connectivity values) [Kelly et al., 2012, Smith et al., 2015].

In our analysis, we focused on four steps involved in processing and post-processing of neuroimaging data. Our results on comparison of the tracking methods demonstrate that connectomes obtained through probabilistic tracking are more consistent than the ones obtained by applying deterministic tracking, supporting previous findings (Fig. 2–3)[Bonilha et al., 2015, Sotiropoulos and Zalesky, 2019].

Although tractography has successfully been applied in noninvasively reconstructing the structural brain network on dMRI data, it is known to generate spurious fibers which introduce direct connections between two regions which would otherwise lack a direct connection [de Reus and van den Heuvel, 2013]. Functional connectomes, on the other hand, is forced to have direct connections between every region pair by design since connectivity is calculated as the correlation between the timeseries of regions, potentially introducing false direct connections. Since false connections are commonly assumed to be among the weak connections, thresholding of weak edges has been a common practice to eliminate noise in the connectomic data. Our results, however, indicate that, thresholding does not alter the consistency of structural connectomes significantly unless edges are heavily pruned (Fig. 4), supporting recent findings [Civier et al., 2019]. This result indicates that, thresholding of weak edges in structural connectomes might not be necessary especially if the analysis involves weighted edges. In thresholding of positive and whole functional connectomes, our results demonstrate that network topology is more consistent than individual connections in response to removing lower weighted edges, while negative connectomes demonstrate a steady decline in consistency with thresholding (Fig. 5). This result might indicate the presence of a steady network with strong edge weights in the positive functional connectomes, while the negative functional connectome lacks such a consistent organisation.

**Figure 4.**
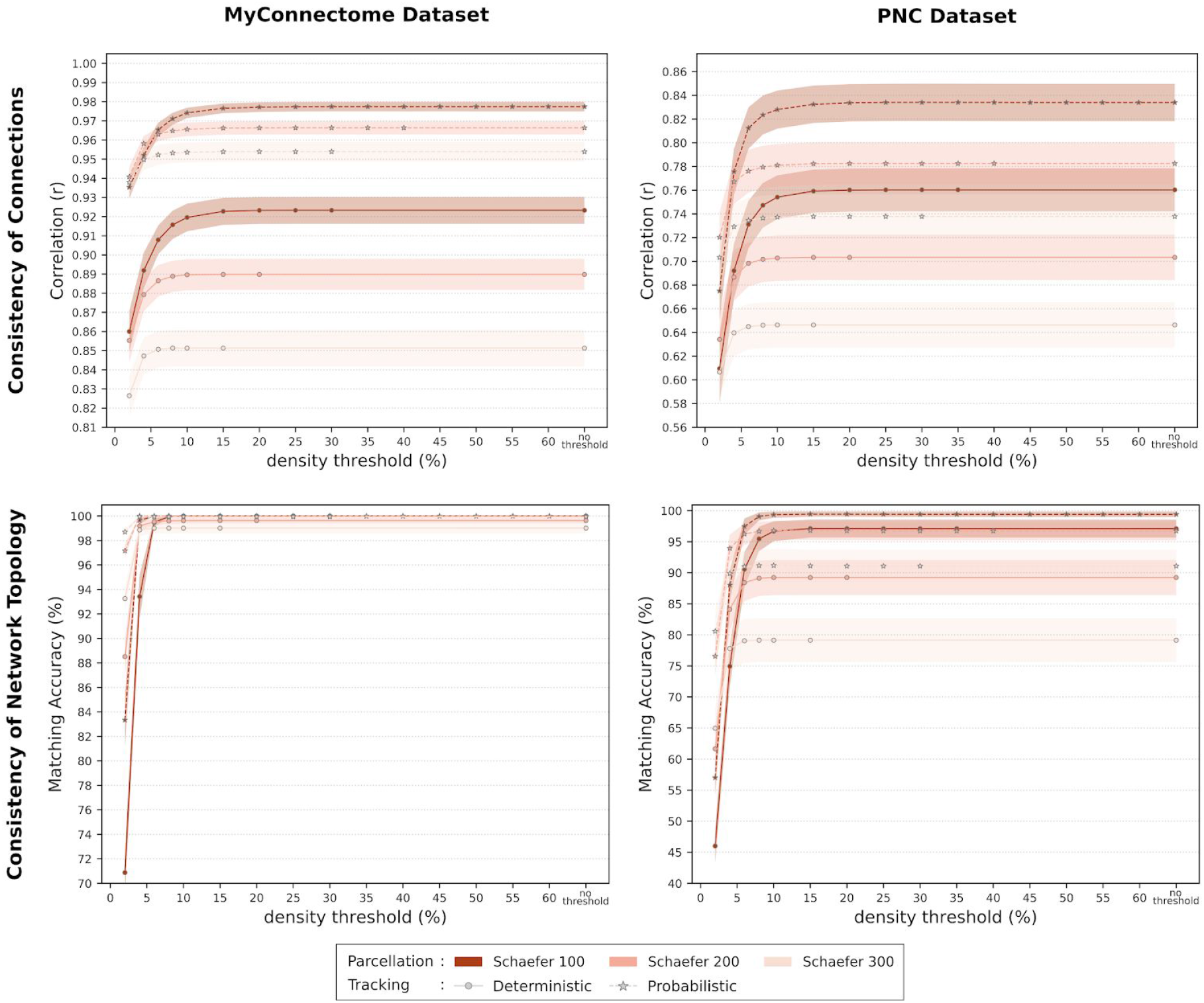
Effect of density thresholding on consistency for deterministic and probabilistic structural connectomes. Top and bottom rows show consistency of connections (correlation) and network topology (matching accuracy), while the left and right columns contrast intra- (MyConnectome) and inter-subject(PNC) consistency, respectively. (Shaded areas indicate standard deviation in consistency scores.)

**Figure 5.**
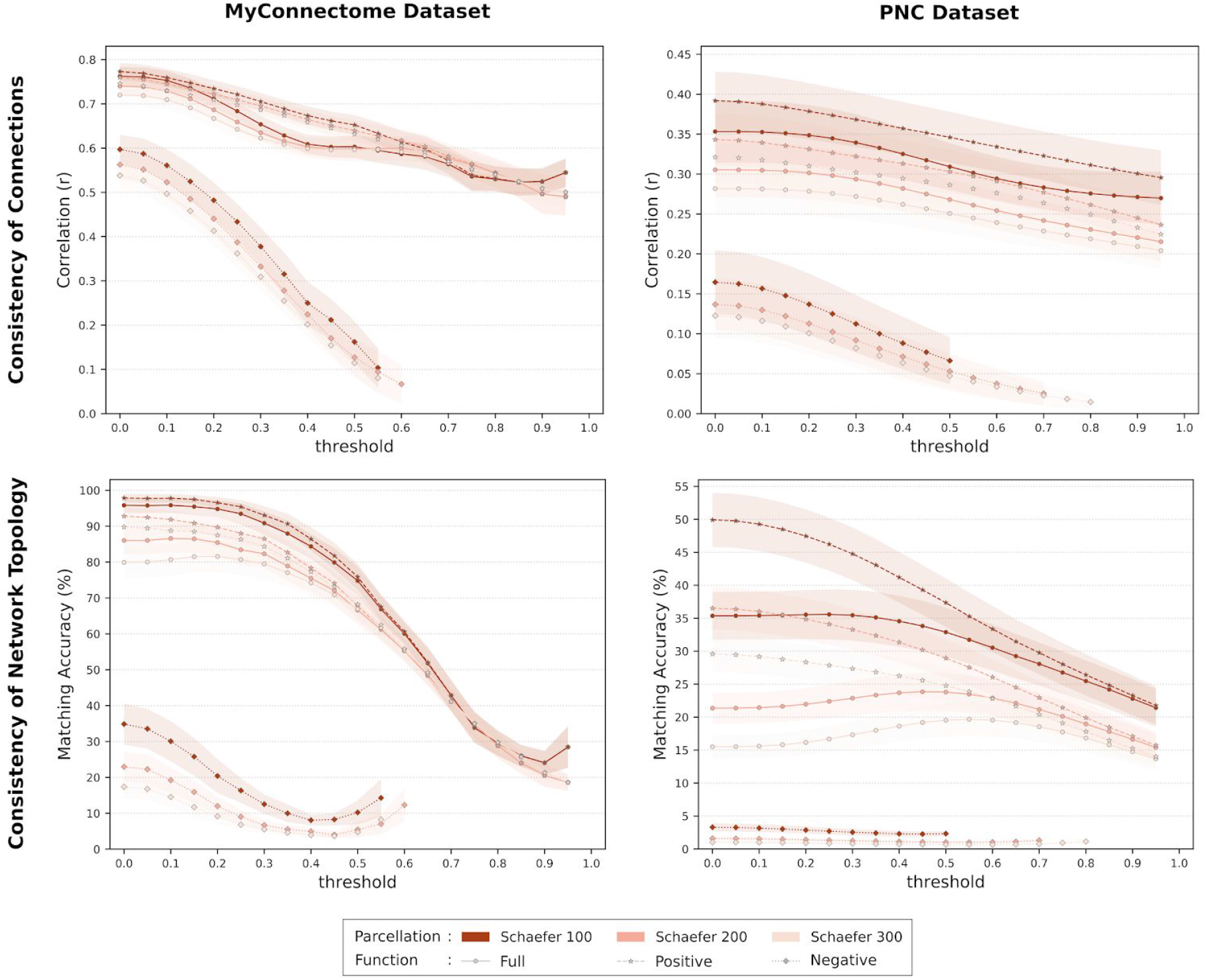
Effect of thresholding on consistency for full, positive, and negative functional connectomes. Top and bottom rows show consistency of connections (correlation) and network topology (matching accuracy), while the left and right columns contrast intra- (MyConnectome) and inter-subject (PNC) consistency, respectively. (Shaded areas indicate standard deviation in consistency scores.)

In analyzing the effect of coarseness of the parcellation on connectomic consistency, we observed that connectomes with lesser number of regions are more consistent within and across subjects in both structure and function, agreeing with the literature (Fig. 1,5) [Zalesky et al., 2010, Cammoun et al., 2012]. This result highlights the importance of the spatial scale of nodal parcellation when making a comparison across studies. Another common post-processing step in generating structural connectomes is in determining the edge weights, which might involve scaling the fiber counts between two regions with the logarithm function or with the total volume of the two regions. Our results indicate that, log scaling increases structural consistency while scaling by node volume does not induce any significant effect (Fig. 2-3). The positive effect of scaling on consistency could be attributed to the fact that log function shrinks the range of edge weights, reducing the variability of connectivity structure across subjects. Thus, although such an effect might be desirable for population studies, it should be avoided in subject specific research.

### Network topology is more consistent than individual connectivities

Human brain is known to have a structural and functional organisation that is consistent across subjects. Although the consistency have been analyzed at the level of individual connections [Guo et al., 2012, Horien et al., 2019] and network topology [Raichle et al., 2001, Owen et al., 2013] separately, a comparison of consistency between these two levels of analysis has been lacking. Our analysis mitigate this need by evaluating consistency at both levels. In the analysis of structural connectome, we observed a near perfect consistency of network topology with MA ranging in [99%,100%], while individual connectivities were farther away from a perfect consistency score as *r* was in [0.85,0.95] range (Fig. 1). A similar pattern was also observed in consistency of functional connectomes. This result indicates that despite variations of individual connections, the overall brain organization is stable within and between subjects. This also demonstrates the effect of similarity measure choice in connectomic consistency analysis as results and ensuing interpretations would differ according to the chosen metric.

### Negative functional connectivity has the lowest consistency among all connectivity types

Although have been available to the neuroscientific community since the beginning of the rs-fMRI studies, the negative functional connectivity is very little understood and its mechanism in the context of network physiology has been a subject of debate. Some studies suggested that it could be an artifact of regression of global signals [Murphy et al., 2009; Weissenbacher et al., 2009] while others demonstrated that it mainly contains long-range connections which might indicate a biological basis [Schwarz and McGonigle, 2011]. Overall, negative functional connectivity is seldom evaluated in neuroscientific research [Chen et al., 2011].

In our analysis, we observed that negative functional connectivity has the lowest consistency among all connectivity types especially with the consistency of network topology being negligible at the PNC dataset (Fig. 5). This can be considered as an evidence against the presence of a biological basis for negative functional connectivity. On the other hand, a decent consistency that we observed on MyConnectome might imply the presence of an underlying biological mechanism. Although inconclusive, our results indicate that systematic longitudinal analyses on negative functional connectivity might provide insights into a possible mechanism behind this phenomenon.

## 5. Conclusion

In this study, we presented a comprehensive analysis of consistency across structural and functional connectomes. We showed that structural connectomes are more consistent relative to functional connectomes, with structural connectomes obtained via probabilistic tracking rendering especially higher and negative functional connectomes demonstrating significantly lower consistency. We also evaluated the effect of some of the connectome processing steps and demonstrated that i) consistency is higher in coarse parcellations, ii) thresholding of weaker edges effect consistency if done heavily, and iii) log scaling of edge weights increases structural consistency. Evaluating a longitudinal and a cross-sectional dataset, we showed that the connectomic consistency of a single subject across time is higher than the consistency across a set of subjects. These results broaden our understanding on the relationship between structure and function of the brain and set a reference point for researchers on connectomic consistency at various setups.

We note that our analysis has certain data related limitations that are challenging to isolate from the study. Firstly, although both datasets are acquired on 3T scanners, difference of acquisition parameters and protocols might have contributed to the consistency variation. Secondly, the age range of subjects in the PNC dataset being [8,22] might have contributed to the lower consistency that was observed as brain is known to have structural and functional changes during development [Satterthwaite et al., 2013, Gu et al., 2015]. Similarly, PNC dataset containing subjects from both sexes might be another factor contributing to lower consistency as structural and functional sex differences were reported in brain connectivity [Ingalhalikar et al., 2014, Tunç et al., 2016]. In future, we will investigate connectomic consistency across development and between sexes.

## Acknowledgments

This work was supported by the National Institutes of Health (NIH 1R01MH117807-01A1).

